# Foundation Model RNAGAN Enhances Biomedical Insight of Nasopharyngeal Carcinoma Metastasis

**DOI:** 10.64898/2026.07.02.736240

**Authors:** Zhaozheng Hou, Yijia Qian, Victor Ho-Fun Lee, Dora Lai-Wan Kwong, Xinyuan Guan, Zhonghua Liu, Wei Dai

## Abstract

RNAGAN (version 2.0, https://github.com/ZhaozhengHou-HKU/RNAGAN-2.0.git) is a published foundation model that analyzes single-cell and bulk-level RNA sequencing samples and enables multiple applications that enhance medical insights. Here we applied this model to Nasopharyngeal Carcinoma (NPC) as in-context few-short format (i.e., the model was never trained with any NPC data). We conducted all four supported functions, which include sample stratification, vectorization, pseudo data generation, and marker identification. The results were then used for identifying metastatic NPC and to investigate mechanisms associated with NPC metastasis.

Examination with stratification showed that the accuracy of RNAGAN results for evaluating the metastasis risk in NPC patients are comparable to or outcompeted recently published risk estimation linear prediction model. Vectorization results present consistency across multiple cohorts and RNAGAN model versions. In the task of identifying markers and mechanisms related to NPC metastasis, incorporating pseudo data substantially enhanced the representativeness of single-cohort-based differential expression (DE) analysis. Moreover, RNAGAN identified metastasis-related marker genes based on single cohort, were concordant with the ground truth obtained across multiple cohorts (p=1.05e-9).

Regarding biomedical mechanisms, RNAGAN enabled second-order feature extraction, unveiling a remarkable domination of the protective function of adaptive immune responses (as indicated by *IL21R* levels) over the hazardous function of chronic, non-resolving innate inflammation (as indicated by *S100A8* levels) against NPC metastasis after first-line treatment. This association demonstrates a high degree of consistency with the external cohort.

This study demonstrates the utility of the foundation model RNAGAN in uncovering therapeutic insights for novel cancer types without extra training. We reveal a critical spatial mechanism preventing distant metastasis via humoral anti-tumor immunity in NPC. High *S100A8* expression by innate antigen-presenting cells (APCs) triggers an inflammatory cascade promoting epithelial-mesenchymal transition (EMT) and metastasis. However, when germinal center *IL21R*+ B cells simultaneously colocalize with these innate signals, they override this suppressive tissue stress. Spatial analysis shows that a high *S100A8*/*IL21R* intersection within tumor regions strictly distinguishes treatment responders, whereas non-responders display spatial mismatch or *S100A8*+ hyper-infiltration. This coordinated innate-adaptive cross-talk sustains functional tertiary lymphoid structures (TLS) that mature IgG-secreting plasma cells, which opsonize and eliminate emerging EMT tumor cells before systemic escape. Consequently, while *S100A8* alone is an unreliable prognosticator, its spatial colocalization with *IL21R* is a robust protective indicator overlooked by conventional bulk analysis methods.

## Introduction

RNAGAN (available at https://github.com/ZhaozhengHou-HKU/RNAGAN-1.0.git) [1] is a published foundation model for biomedicine research, designed with generate adversarial network (GAN) structures [2, 3]. This model aims to process RNA sequencing data at single-cell or bulk level and is trained with real human samples only. The major advantages include the capability with small datasets (20 to 30 samples), its explicit pathway enrichment analysis layer and the corresponding feature extraction, and the structural prevention of data memorization and privacy leakage [1, 4].

Same as other foundation models that have demonstrated remarkable capacity for biomarker discovery and patient stratification [5], the clinical and biomedical applications of these models are hindered by the lack of comprehensive and detailed validation for diseases or cancer types that fall outside of the training distributions [6, 7]. While common malignancies are often well-represented in large-scale repositories, rare or regional diseases or cancer types lack the necessary longitudinal data and molecular profiling consistency required to rigorously test model generalizability [8, 9]. For this consideration, the evaluation with a “Goldilocks” selection of cancer type for evaluation, sufficiently rare to ensure genuine out-of-distribution evaluation, yet have widely accepted biomedical insights and enough standardized molecular depth to permit statistically significant benchmarking is of particular value [10].

In this study, we chose Nasopharyngeal Carcinoma (NPC), particularly its metastasis and response to treatments as the tasks for evaluating the performance of RNAGAN. NPC is a malignancy occurring at nasopharynx, with a high incidence in southern China, Southeast Asia, and parts of North Africa [11–13]. The development of clinical oncology in recent decades have confirmed the importance of tumor microenvironment (TME) to the development, metastasis and drug resistance [14, 15]. The TME is an aggregation of stromal, immune, tumoral cells, and extracellular components within tumorous regions. It is also where mechanisms such as adaptive immune responses [16] and nonspecific inflammation [17] determine cancer outcomes, and so as the tertiary lymphoid structures (TLS) formed to promote apoptosis of malignant cells and enhance immunotherapy response [18]. Despite the fact that research on these metastasis-related mechanisms has been conducted for decades, as is the case with other cancer types, there are still many details that remain unknown, particularly the interactions and dominations of these mechanisms.

In this study, we evaluated the benefit of RNAGAN in NPC metastasis-prediction and related tasks with two bulk RNA sequencing data sets from Hong Kong and Guangzhou [19], one single cell data set [19], and one spatial transcriptomic data set [18]. We conducted all four supported functions, which include sample stratification, vectorization, pseudo data generation, and marker identification. Particularly, one of the second order features (association of two marker genes) related to distant metastasis was evaluated in detail. Moreover, we introduced two more updated AI reasoning mechanisms to RNAGAN and established enhanced versions (RNAGAN-Quadra and RNAGAN-GPT).

## Materials and methods

### Data collection and study approval

All the data sets previously employed for training the original RNAGAN [1] have also been included in this study and utilized for the training of RNAGAN version 2. These include 14 single-cell RNA-seq datasets, comprising a total of 4.6 million cells, and 8 bulk RNA-seq datasets, encompassing 5,900 samples. These data were obtained from the CZ CELLxGENE Discover [20], the Cancer Genome Atlas (TCGA) database [21] and the Gene Expression Omnibus (GEO) [22]. Further information can be found in the supplementary document of the paper that released RNAGAN [1].

Regarding the NPC samples, we employed bulk-sequencing data from two cohorts, one from Hong Kong and Guangzhou correspondingly. Hong Kong cohort (cohort 1), with ethics approval UW 24-041 and UW 09-125, contains 77 NPC patients at four Hong Kong hospitals; and the other from Guangzhou (cohort 2) with ethics committee approval G2021-110-01, contains 100 patients at Sun Yat-sen University Cancer Center. Details of these cohorts can be found in previously published study [19]. To validate the single-cell level performance of RNAGAN, we used the single cell data collected from ten Guangzhou NPC patients. For further details, refer to the corresponding study by Liu et al. [23]. Doublets and proliferating cells were removed using the same procedure for data preparation of the signature matrix generation in previous study [19].

### New developments in RNAGAN 2.0

In accordance with the published RNAGAN version 1.0 [1], two significant modifications have been implemented with the objective of enhancing the system’s robustness and mechanical support of biomedical features: the replacement of the normalization layer, and the introduction of two variations of representation shaping layers at the network bottleneck (details can be found in **Figure S1**).

RNAGAN 1.0 adopted batch normalization (normalization of layer outputs across all input cases within a single batch during network training), and they are replaced by layer normalization [24] (normalize the layer outputs for each input case) in the new version. This replacement is for the consideration that the network should facilitate stable process and reasoning for each input case, regardless of the condition of other inputs that provided in the same batch (which could originate from different cohorts in network training, and not available in application scenarios). As indicated by prior research, this replacement has the potential to enhance both the precision and the extent to which the results generalize [25, 26]. Notably, this modification does not encompass the final normalization layer for discriminators. The rationale behind this choice stems from the fact that the training procedure for discriminators consistently utilizes balanced positive and negative input cases for each batch, an approach that has been recommended by numerous studies [27].

For bottleneck representation shaping, we introduced two variations to replace the initial fully-connection layer: The first variation is the quadratic layer (designated as RNAGAN-Quadra in this paper) [28, 29], and the second variation is the transformer block [30, 31] (designated as RNAGAN-GPT). The quadratic layer is designed to compute the squares and product values of the input for this layer, thereby ensuring that the downstream process can directly base its operations on the values such as *x*_1_^2^ and *x*_1_ ∗ *x*_2_. This design explicitly supports the understanding of U/J-shaped non-linearity and interactions (epistasis) between genes and/or pathways, which are extremely common in biomedical mechanisms [32]. Multiple authoritative studies and practical experiences have reported the performance improvement of this design [26, 33]. The transformer block has garnered a more substantial degree of recognition, and hybrid U-net-like architectures with the transformer serving as the representation shaping layer have demonstrated their efficacy in numerous recent studies [34, 35].

The training of RNAGAN version 2.0 utilized the same loss functions, data sets, and settings as version 1.0 [1]. This process is comprised of three steps including (1) initial training of the discriminator for cell type identification; (2) fine training of both the generator and the discriminator for pseudo single-cell data generation and cell type identification; and (3) transfer training of both the generator and the discriminator for pseudo bulk-sequencing data generation and patient stratification.

### Function 1: Stratification

In this study, we carried out stratification for single-cell (with the task of identifying tumor cells), and for bulk-sequencing data (with the task of identifying NPC patients that will develop distance metastasis within 3 years after first-line treatment). Single-cell task was accomplished in the positive reference only format, because the cells are clustered into 10 types [19], and 20 randomly selected negative samples may not be representative enough. Bulk-sequencing validation used both positive (metastatic NPC) reference group and negative (non-metastatic NPC) group.

The function of RNAGAN’s stratification is achieved by its discriminator sub-network, which is designed to evaluate the similarity of one given sample to the reference sample group. The stratification can be achieved through the provision of a positive reference group alone or a combination of a positive and negative reference group, as illustrated below:

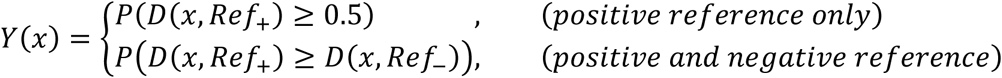

RNAGAN 2.0 employed 20 samples to establish the reference group for each time, and the probability can be evaluated using the bootstrap method.

### Function 2: Vectorization of sample populations

Apart from the metrics directly evaluating the stratification accuracy, vectorization results is also an approach to confirm the representation validity / biological fidelity of RNAGAN. A foundation model that captures correct and biologically meaningful features, should present the division of different patient groups (such as metastatic and non-metastatic NPC), and this pattern should be robust across different cohorts. For both cohort 1 and cohort 2, 100 vectors were generated for metastatic/non-metastatic NPC populations using the bootstrapping method. The principal component analysis (PCA) was computed for the purpose of visualization.

The generator sub-network of RNAGAN can generate a 64-dimentional vector for a given group of 20 samples as reference. The generated vector is for the given reference group rather than one single sample, because the variance between references is also incorporated into the vectorization procedure.

### Function 3: Pseudo Data Generation

Pseudo data can be used to improve the representativeness and robustness of sample-based downstream analysis, such as differential expression (DE) analysis. In this study, we generated the pseudo data for cohort 1 for the metastatic/non-metastatic NPC population by sampling 20 samples for each time from the corresponding population as the reference group. This procedure generates one pseudo sample at a time and was repeated multiple times to generate a pseudo population with a 1:1 sample size ratio as cohort 1. Pseudo data was incorporated into Cohort 1, with the ratio ranging from 0% (real data only) to 100% (half real, half pseudo data). The up- or down-regulated marker genes were subsequently filtered using the Mann-Whitney

U test, with a p-value less than 0.01. With respect to the ground truth for performance evaluation, genes that demonstrated significant up- or down-regulation in the metastatic NPC population in both cohort 1 and cohort 2 (Mann-Whitney p ≤ 0.05 in both cohorts) were designated as the ground truth.

To elaborate, the objective of this section of validation is to ascertain whether RNAGAN can facilitate the generation of a gene list exclusively based on cohort 1. The primary outcome of this process is expected to be concordant with the intersection of cohort 1 and cohort 2, as this combination is regarded as a more authentic representation of the underlying NPC-related mechanisms and markers.

### Function 4: Gene and Pathway Marker Identification

We extracted the marker genes and pathways that related to NPC metastasis risk, with the aim of interoperating the stratification results from RNAGAN and also identify markable features and mechanisms to gain biomedical insights of NPC metastasis.

The feature extraction function of RNAGAN has the capacity to generate a list of metrics for marker identification based on positive and negative samples that have been provided:

i. **Classic statistics**: Fold-change, p and adjusted-p from Mann Whitney U test;
ii. **First order network features**: Feature-output gradient, Grand CAM scores; (iii)**Second order network features**: Second order Feature-output gradient, and R2 scores (coefficient of determination).

The first and second order gradients are defined as follows:

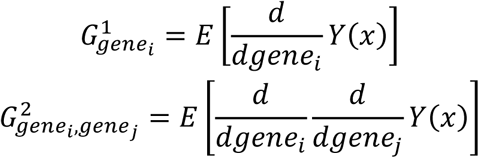

For both gene and pathways, the standards for marker identification are as follows: Absolute log2 fold change >= 1, Mann Whitney p <=0.05, and averaged first order gradient above or lower than 0 (consistent with the log2 fold change).

For identified markers, the second-order association between these markers with R2>=0.05 are considered as significant associations.

### Spatial Transcriptomic Analysis

To validate the identified gene-level association between *S100A8* and *IL21R*, we adopted FFPE Visium data from 12 patients in the study by Liu et al.[18]. These patients received toripalimab plus chemotherapy and were assessed as responders/non-responders based on the RECIST v.1.1 criteria [36]. The metric Intersection over Union (IoU) was adopted to quantify the spot-level colocalization of these two marker genes, which is defined as follows:

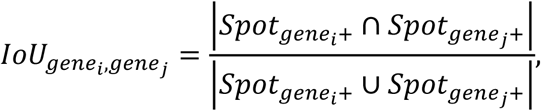

where *Spot_gene_i_+_* refers to the Visium spots with SCTransform values larger than 1.

## Results

We conducted all four functions as stated in the introduction paper of RNAGAN [1], which include: sample stratification, marker identification, pseudo data generation, and vectorization. Subsequently, the results were utilized to investigate underlying mechanisms associated with NPC metastasis.

### RNAGAN can Estimate NPC Metastasis Tendency with Clinically Usable Accuracy and Biological Fidelity

In this study, we applied the AI model to identify patients developed distant metastasis within three years after the first-line treatment (according to the RNA-sequencing results of the primary tumor biopsy before the first-line treatment) (**Figure 1A-B**) It is noteworthy that all of these results are derived from in-context few-short applications, and the model has not been trained on any NPC data.

**Figure 1.**
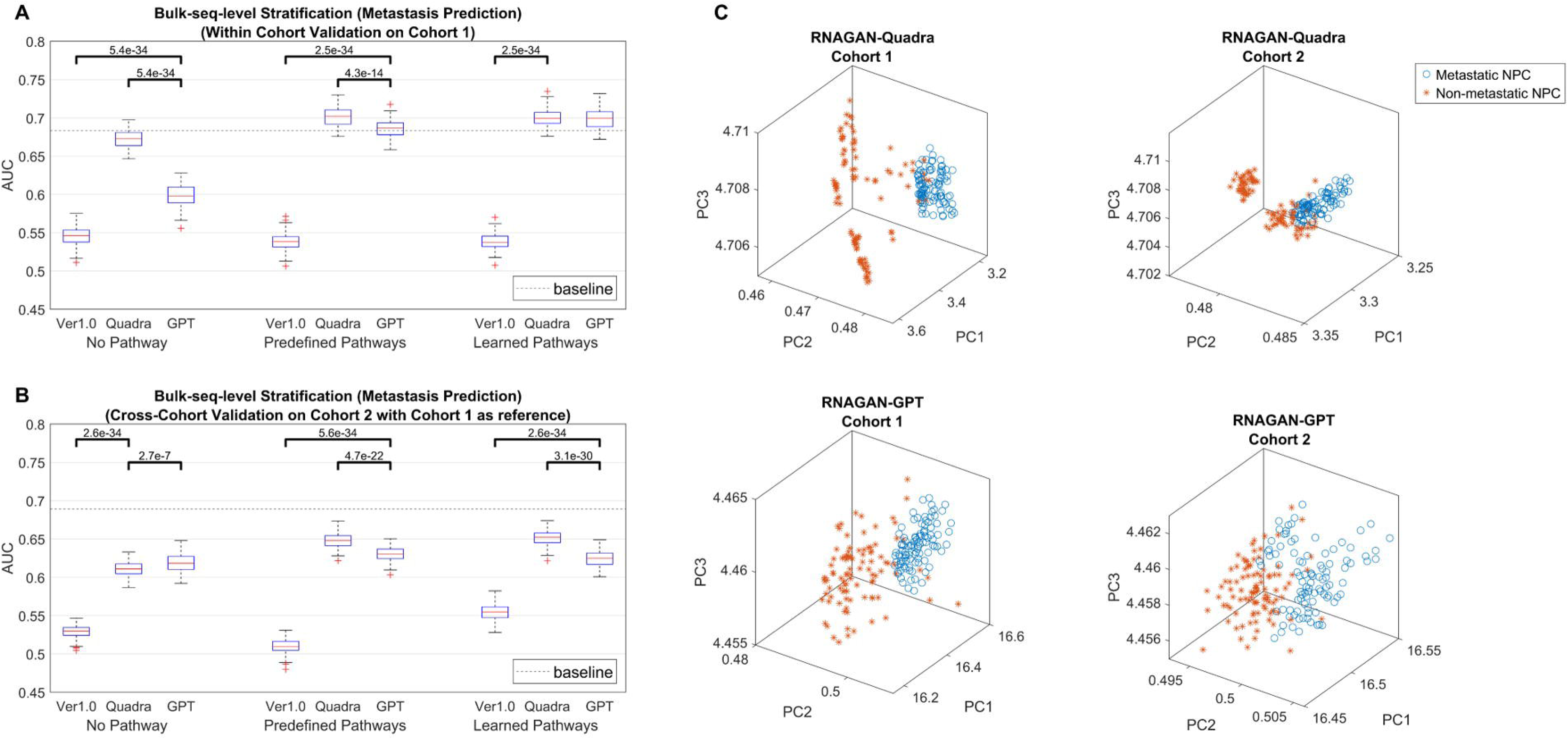
Stratification from RNAGAN and corresponding vectorization results. **(A)** Identify metastatic NPC within cohort 1, both predefined pathway versions and learned pathway versions achieved AUC that is comparable with or even better than the baseline (a recently published linear model for stratification). **(B)** Cross cohort application of metastases risk estimation, the performance is significantly worse than within cohort application situations, and it does not outcompete the baseline (but notice that RNAGAN is not designed or trained for cross-cohort application). **(C)** Two independent cohort present consistency in the latent space of both RNAGAN Quadra and GPT versions. Moreover, although Quadra and GPT versions have never interacted with each other during the training, the vectorizations in PCA show similar spatial locations.

RNAGAN 2.0 (both Quadra and GPT versions) achieved significantly better performance than RNAGAN 1.0. Particularly, the Quadra versions achieved the accuracy level that is comparable to or even outcompeted a recently published linear prediction model [19] (derived from typical clinical medicine research (LASSO) [37, 38]) in the within cohort test. However, the performance significantly reduced in the cross-cohort application test (building reference groups with cohort 1 samples and evaluate metastatic risk for each NPC sample from cohort 2). The versions with predefined pathways (reported gene sets from Human Molecular Signatures Database as detailed in [1]) led to higher AUCs compared to the versions without the pathway layer. However, the versions with learned pathways (same number of pathways but the weight factors were learned from scratch based on during single-cell data training) did not present a significant improvement in the accuracy. Therefore, it is reasonable to conclude that the most recommended version for metastasis risk stratification is RNAGAN-Quadra with predefined pathways.

We generated the vectorization results for metastatic and non-metastatic NPC populations from the latent spaces of RNAGAN Quadra and GPT versions (**Figure 1C**). The vectors demonstrate a significant separation between NPC patients with different clinical outcomes, and the spatial relationship in the latent space is consistent across cohorts 1 and 2. Furthermore, despite never interacting with each other during training, the Quadra and GPT versions exhibit similar spatial location patterns in PCA. The PC1 for RNAGAN-GPT is flipped to show consistency, which is reasonable since PCA only considers the axes of principal components (PC), and the positive direction is random and mathematically equivalent.

We have also evaluated the performance of RNAGAN with few other tasks: single-cell-level application that identify NPC tumor cells from other cell types, achieving area under the curve (AUC) scores above 95%; and the stratification of three molecular subtypes [39], with the AUC values exceeding 80% for within-cohort applications (**Figure S2**).

### RNAGAN Improved the Representativeness Differential Expression Analysis

Pseudo bulk-sequencing data was generated based on cohort 1 and conducted a differential expression (DE) analysis of metastasis-related marker genes with the data mixed with real and pseudo data with different amounts of pseudo data (**Figure 2**).

**Figure 2.**
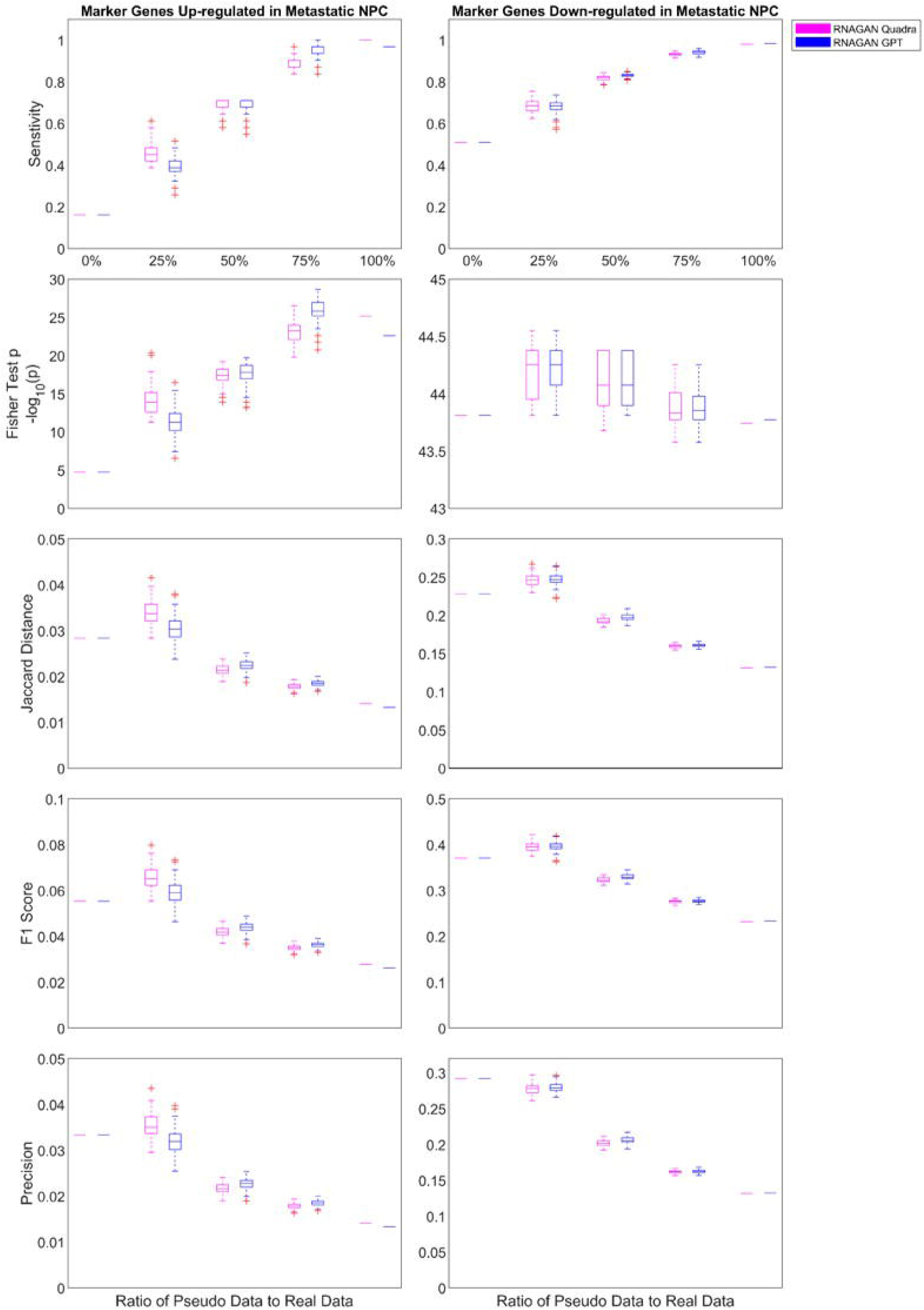
Metrics of evaluating the benefit of pseudo data for as differential expression (DE) analysis of metastasis-related marker genes.

RNAGAN Quadra and GPT versions present very similar trends of metrics regarding different ratios of pseudo data to real data. The sensitivity of DE analysis kept improving by adding more pseudo data and achieved 95% sensitivity with 1:1 pseudo-real data ratio from less than 60% by real data only. It means that RNAGAN generators captured more than 95% of “real” marker genes and helped to make these markers more significant with the generated pseudo data. The p values from fisher exact test for up-regulated markers improved with more pseudo data, suggesting that these generated data improved the enrichment of real markers in the DE analysis (the improvement in down-regulated markers is not as significant because the p value is already very significant from cohort 1 by its own). However, F1 score and precision decreased since adding more than 25% of pseudo data, suggesting that RNAGAN was also adding its own features to the data and introduced some false-positive markers.

In conclusion, adding 75-100% pseudo data helped to reduce type II errors (false negatives that miss true markers). If users also want to avoid type I errors (false positives that introduce false markers into the downstream analysis), adding 25% pseudo data significantly improves marker filtering. It is important to note that these evaluation metrics (such as Wilcoxon p values) would not improve by simply repeating the real samples. This indicates that RNAGAN truly makes novel contributions to the analysis and leads to more robust and representative results.

### First and Second Order Metastasis-related Features Identified by RNAGAN

We extracted and visualized gene and pathway level features related to NPC metastasis base on cohort 1 data (**Figure 3**). Since RNAGAN-Quadra with predefined pathway achieved highest AUC in the task of metastasis estimation, this version was employed to extract the features.

**Figure 3.**
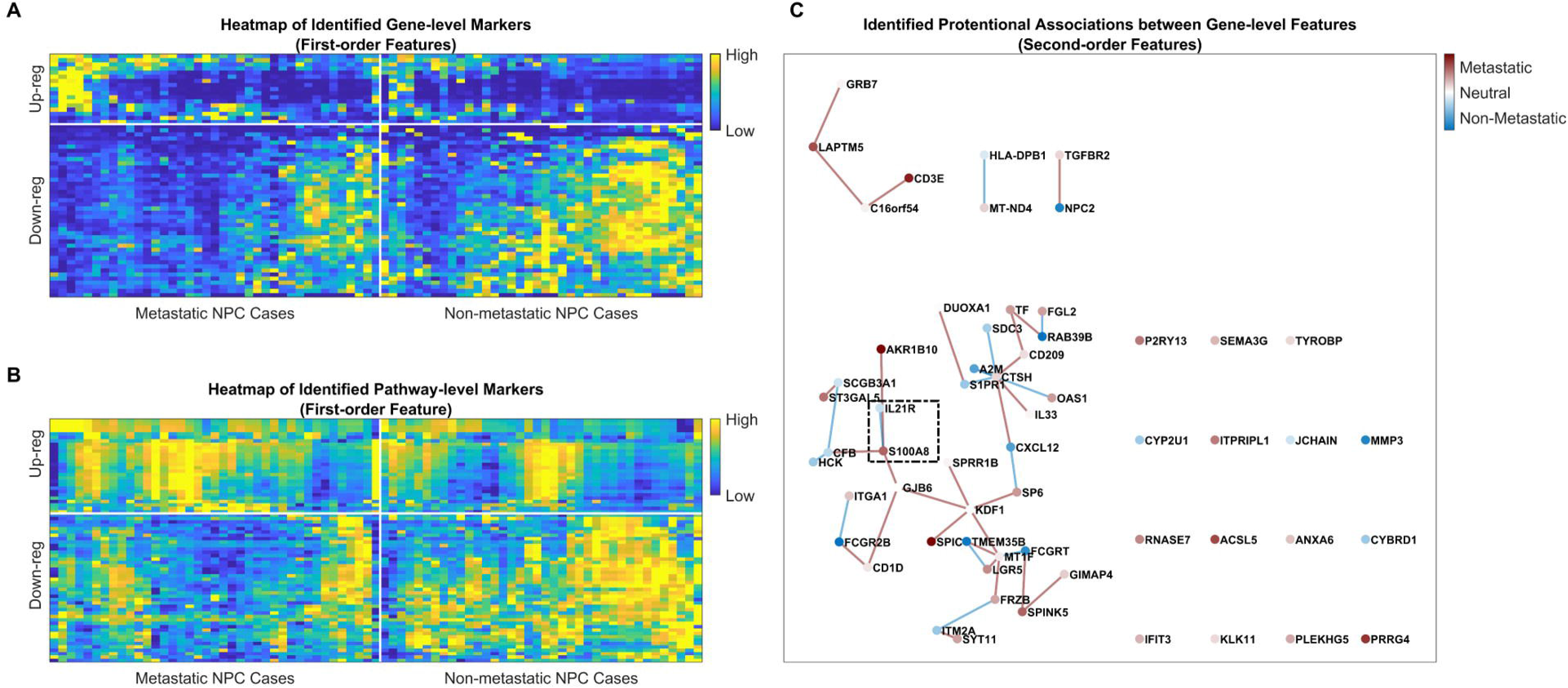
Identified first and second order features of NPC metastasis. Features and the associations were generated with RNAGAN-Quadra with predefined pathway, and base on Cohort 1. **(A)** Heatmap of the expression level of identified marker genes. **(B)** Enrichment of identified marker pathways. **(C)** Identified associations between marker genes. The distance between connected genes were decided by the significance level of the associations (closer genes have more significant associations).

The network identified 59 marker genes and 98 pathways that make significant positive contribution to correct estimations (**Table S1**). The identified metastasis-related marker genes were concordant with the ground truth obtained across cohort 1 and cohort 2 (Fisher exact p=1.05e-9). RNAGAN can also identify second-order features (associations between markers rather than hazardous/protective effects represented by single markers). The mechanisms suggested by the corresponding plot (**Figure 3C** for marker genes, **Figure S3** for marker pathways) can be interpreted with the table as follows:

**Table 1.** Interpretation of identified second-order features.

| Feature A | Feature B | Connection | Suggested Association |
| --- | --- | --- | --- |
| Hazardous+ | Hazardous+ | Hazardous+ | Synergistic hazardous effect. |
| Hazardous+ | Hazardous+ | Protective- | Non-additive interaction or ceiling effect. |
| Hazardous+ | Protective- | Hazardous+ | Feature B is protective, but hazardous feature A dominates the outcome. |
| Hazardous+ | Protective- | Hazardous- | Feature A is hazardous, but protective feature B dominates the outcome. |

We highlighted an association of *S100A8* and *IL21R* as an example. The second-order feature analysis indicated that *S100A8* represents a hazardous mechanism, and a protective mechanism represented by *IL21R* dominates the metastasis outcome of NPC patients. We further investigated this mechanism with spatial transcriptomic data in later sections.

### Identified Association between *S100A8* and *IL21R* Validated in Multi-omics Datasets

From the gene-level associations identified by RNAGAN, we chose the domination of *IL21R* levels over the hazardous function indicated by *S100A8* levels as an example to carry out further investigation (**Figure 4**).

**Figure 4.**
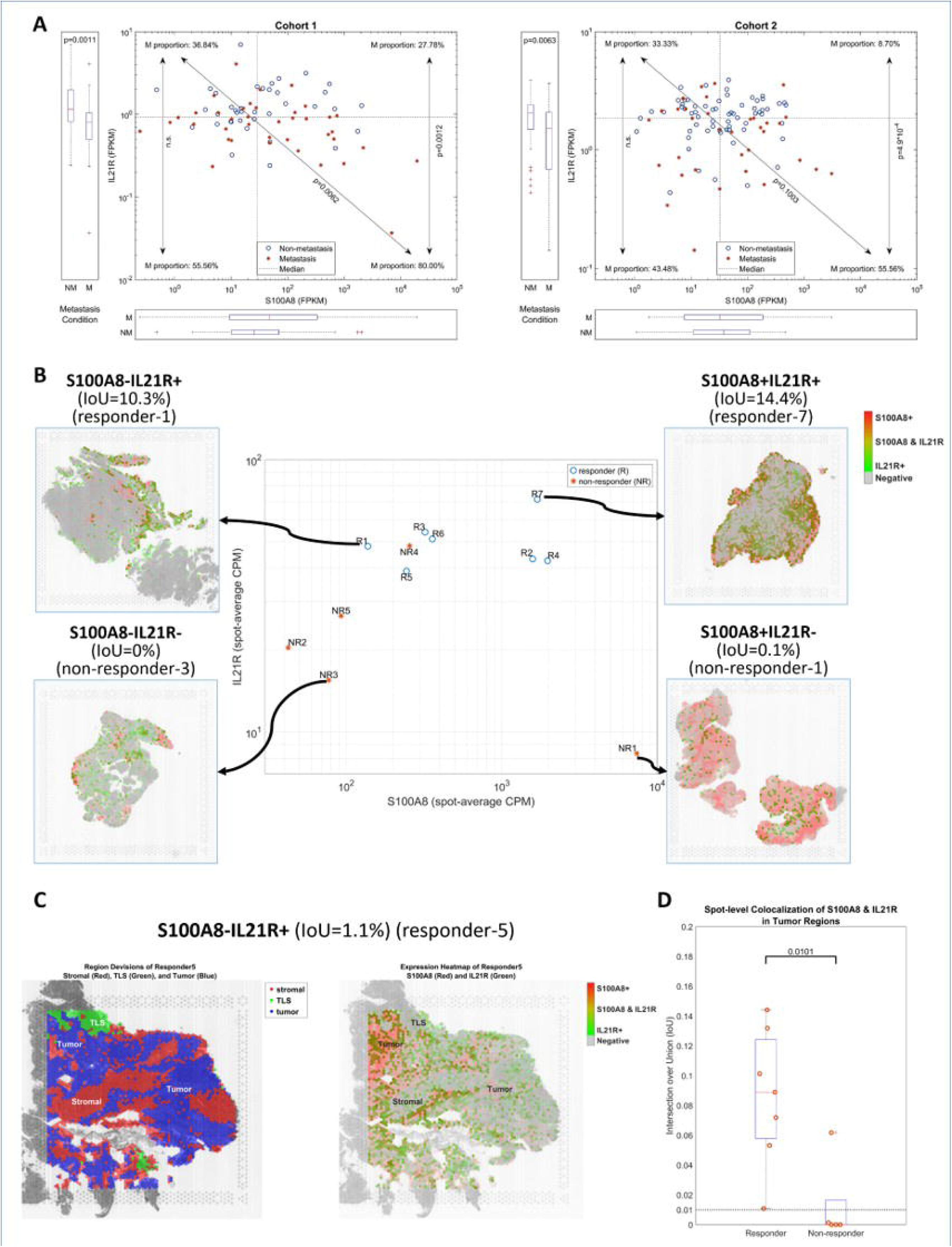
Validation of the role of *IL21R* and *S100A8* in the NPC’s response of treatment. (**A**) *IL21R*+ patients are less likely to develop distance metastasis after receiving the first line treatment, and this difference becomes particularly significant in *S100A8*+ patients. For *IL21R*-patients, higher *S100A8* indicates higher risk of metastasis. Although this trend and association is identified by RNAGAN base on cohort 1 only, same pattern can be observed in NPC cohort 2. (**B**) Sample-level analysis of spatial transcriptomic dataset also presented similar patterns observed in bulk-sequencing data set, and *IL21R* shows enrichments in *S100A8*+ areas in the samples with high *S100A8* expression level. (**C**) Spatial sequencing data from responder 5 shows the representative spatial locating pattern of *S100A8* and *IL21R*: *S100A8* are highly expressed in tumor regions and not expressed in TLS region, it can also be found in stromal cell region particularly in non-responders; *IL21R* is expressed in all three regions and shows colocalization with *S100A8* particularly in *S100A8*+ samples. (**D**) Responders have significantly higher IoU scores of *S100A8* and *IL21R* in tumor regions, suggesting the significant role of colocalization of these two genes.

As indicated by RNAGAN, *IL21R*+ cases are less likely to develop distant metastasis after receiving first-line treatment, while high *S100A8* expression indicates a higher metastasis risk, and the effect of *IL21R* dominates the overall risk level over the mechanism represented by *S100A8*. In other words, *S100A8*+*IL21R*+ cases are more likely to respond well to treatment, while *S100A8*+*IL21R*-cases have a worse response.

Although RNAGAN identified the above association based on cohort 1 data only, this pattern was also observed in cohort 2 (**Figure 4A**) and so as spatial transcriptomic dataset in terms of the response to immunotherapy (**Figure 4B**). A detailed view of the representative slides in **Figure 4B** illustrates how the spatial organization of innate antigen-presenting cells (APCs) shapes the adaptive germinal center response. Non-responder 3 entirely lacks overlapping signals for the two marker genes, suggesting that the recruitment and activation of adaptive, germinal center *IL21R*+ B cells are not initiated when the baseline stimulus from innate *S100A8*+ APCs (such as monocytes, dendritic cells, or macrophages) falls below a critical threshold. Conversely, responder 1 and 7 (as examples) exhibit robust recruitment of germinal center *IL21R*+ B cells, demonstrating that even a modest presence of *S100A8*+ APCs can successfully drive adaptive humoral immunity. Non-responder 1, however, highlights a dysregulation in this cross-talk that *S100A8*+ APCs widely infiltrate the stromal regions and dominate the tissue landscape, yet germinal center *IL21R*+ B cells remain scarce and isolated, indicating that overabundant, hyper-inflammatory innate myeloid cell accumulation correlates with a failure to sustain the structured microenvironment required for localized germinal center reactions.

Beyond these broad trends, we observed more nuanced spatial distribution patterns across our dataset, as demonstrated by the representative architecture of responder 5 (**Figure 4C**). Here, *S100A8* expression is highly localized within tumor regions but completely excluded from tertiary lymphoid structures (TLS). Notably, *S100A8*+ APC presence extends into stromal cell regions, predominantly in cases characterized by clinical non-response to treatment. In contrast, *IL21R*+ germinal center B cells populate all three microenvironmental regions (tumor, stroma, and TLS) and display pronounced colocalization with *S100A8*+ cells, particularly within *S100A8*-enriched samples.

To quantify this cellular interplay, we compared the Intersection over Union (IoU) of *S100A8* and *IL21R* signals within tumor regions (**Figure 4D**). This spatial metric revealed that good responders to therapy exhibit significantly higher IoU values than non-responders. Critically, 100% of responders maintain an IoU above a minimum threshold of 0.01, whereas 80% of non-responders fall below this baseline. Our spatial analysis indicates that low IoU values stem primarily from two distinct structural pathologies: either adaptive *IL21R*+ cells fail to enrich within *S100A8*+ tumor zones, or the tumor regions become almost entirely saturated by *S100A8*+ APCs, effectively crowding out adaptive immune responders.

Importantly, while elevated *S100A8* expression contextually amplifies the prognostic power of *IL21R*, rendering it a more substantial protective marker, *S100A8* expression alone is not a robust predictor of clinical outcome. Consequently, standard bulk or non-spatial analytical methods are highly likely to overlook this coordinated protective mechanism, which we found to be consistently reproducible across multiple independent patient cohorts through RANGAN.

## Discussions

In this study, we applied the RNAGAN (a published AI foundation model for analyzing single-cell and bulk-level RNA sequencing data) to Nasopharyngeal Carcinoma, a cancer type that has not been included in the training data set of this tool. All the four suggested analysis functions have been carried out and validated in the in-context few-shot mode (i.e., directly use the model to process NPC sample data without model fine-tune): sample stratification, vectorization, pseudo data generation, and marker identification. In addition, we have selected one of the mechanisms identified by this model and validated with an independent cohort and spatial transcriptomic dataset.

Regarding the task of identifying metastatic NPC, RNAGAN has demonstrated the capacity to distinguish metastatic NPC patients with AUC scores that are comparable to or exceed those of a recently published linear estimation model. And vectorization of metastatic and non-metastatic NPC presented a high degree of consistency in the latent representations (with PCA) across both RNAGAN versions and cohorts, suggesting that key underlying structures of the data have been robustly captured and processed.

In terms of identifying metastasis-related markers and mechanisms, adding pseudo data from RNAGAN into real data sets, made the downstream differential expression (DE) analysis more robust. The sensitivity of ground truth gene markers exhibited an increase from less than 60% to over 95%. The F1 score significantly improved and maximized by adding 25% of pseudo data. RNAGAN identified 59 marker genes and 98 pathways that contribute to correct estimation of NPC metastasis risk. It also identified 41 associations between genes, and 37 pathway associations.

A major challenge in contemporary cancer transcriptomics is the reliance on single-marker prognostic indices, which frequently oversimplify the complex, multicellular dynamics of the tumor microenvironment (TME). In this study, our foundation model, RNAGAN, bypassed these classical constraints to uncover a novel, highly coordinated protective mechanism governed by the spatial interplay between *S100A8* and *IL21R*. Remarkably, while this gene-level association was initially identified within our primary cohort, its distinct distribution pattern was independently validated across both an external bulk-sequencing cohort and a spatial transcriptomics dataset.

The clinical significance of this relationship lies entirely within its spatial coordination, a feature that standard analytical pipelines routinely overlook. When analyzed in isolation, *S100A8* does not emerge as a statistically robust or salient indicator of patient survival, leading to its frequent dismissal in conventional bioinformatic workflows as irrelevant noise. However, by evaluating the spatial and contextual dependencies between markers, our AI-driven approach demonstrated that high *S100A8* expression serves as a critical prerequisite that amplifies the protective value of *IL21R*. This highlights the indispensable role of advanced RNAGAN model in discovering non-linear, multi-cellular interactions across independent cohorts that are otherwise invisible to traditional bulk-sequencing analyses.

At the mechanistic level, this interaction reflects a finely balanced crosstalk between innate inflammation and adaptive humoral immunity, which dictates whether a tumor undergoes metastatic dissemination or localized eradication. High baseline expression of the innate alarmin *S100A8*, primarily secreted by monocytes, dendritic cells, and macrophages, triggers an upstream inflammatory cascade mediated via the *TLR4* receptor and the *NLRP3* inflammasome. This pathway accelerates downstream IL-1 signaling, a known driver of tumor progression that promotes epithelial-mesenchymal transition (EMT) and empowers NPC cells to acquire motile, invasive properties.

Our quantitative spatial analysis further refines this model by demonstrating that the physical architecture of these cells directly correlates with therapeutic success. As evidenced by our Intersection over Union (IoU) metrics, a definitive hallmark of clinical responders is a high spatial overlap between *S100A8* and *IL21R* signals within the tumor zones. This spatial proximity ensures that the newly organized TLS can efficiently orchestrate the local maturation of naïve B cells into IgG-secreting plasma cells. Situated at the invasive tumor border, these plasma cells immediately opsonize emerging EMT tumor cells, marking them for antibody-dependent cellular cytotoxicity (ADCC) or phagocytosis before they can successfully intravasate and escape into the systemic circulation.

Conversely, clinical non-responders primarily present with two spatial pathologies that yield a low IoU, indicating a complete failure of *IL21R+* B cells to enrich within *S100A8+* inflammatory zones, or a hyper-dense infiltration where *S100A8+* myeloid cells entirely saturate the tumor stroma. In the latter scenario, the overabundant innate population effectively crowds out adaptive immune responders, disrupting the microenvironmental formatting required to sustain TLS architecture. Taken together, these findings prove that *S100A8* is not a monolithic “hazardous” factor; rather, its clinical outcome is entirely contextual. When spatially synchronized with *IL21R+* B cells, it acts as the initial innate spark that ignites a protective, localized humoral response capable of neutralizing metastatic progression.

In summary, this work demonstrates the significant value of the foundation model RNAGAN in establishing robust medical applications and insights for cancer types it encounters for the first time without extra training. This study enhances confidence in the implementation of this model for other cancer types and diseases.

### Future Prospective

A notable benefit of RNAGAN is its capacity to expeditiously establish the stratification of samples with designated positive and negative references. This process occurs without requiring the model to be informed of the standard or the mechanism that differentiates the reference groups. The foundation model has the capacity to analyze gene and pathway-level markers, as well as the potential associations between them. Consequently, in the subsequent phase, we will employ RNAGAN to conduct a more comprehensive investigation of the markers and mechanisms associated with the various NPC molecular subtypes and clinical outcomes, including relapse.

As a foundation model, RNGAN has been demonstrated to have structural compatibility with virtual gene perturbation; further validation with multiple datasets is currently underway to ascertain the corresponding functionality.

## Author contributions

WD supervised this study. ZH and YQ contributed to data acquisition. ZH developed the study methodology and contributed to data analysis and interpretation. WD and ZH acquired the funding for this project. All authors contributed to writing, reviewing, and revising the manuscript.

## Acknowledgements

This work was supported by the Hong Kong Health and Medical Research Fund (11221746) (to ZH) and Theme-based Research Scheme (T12-703/22-R; T123-70323-N) from the Research Grant Council (to WD).

## Data Sharing Statement

RNAGAN version 2.0, the trained network and related software scripts are available on https://github.com/ZhaozhengHou-HKU/RNAGAN-2.0.git.

## Supplementary figures and tables

**Figure S1**. Representation shaping structures of different RNAGAN versions. **(left)** RNAGAN 1.0 processed the data with FC and leaky-ReLu layers; **(middle)** RNAGAN-Quadra introduced a quadratic layer to capture second-order mechanisms; **(right)** RNAGAN-GPT processed the data with a transformer block.

**Figure S2**. Stratification results from RNAGAN. **(A)** In the task of identifying NPC tumor cells, RNAGAN-Quadra and RNAGAN-GPT significantly outperformed RNAGAN 1.0, and achieved AUC above 90% without pathway layer, and above 95% with the pathway layer, however, learned pathway versions are not as accurate (completely learned by AI from single-cell data training). **(B)** Identification of molecular subtypes of NPC patients. Both Quadra and GPT versions achieved AUC higher than 80% in the within-cohort applications, and performance worse in cross-cohort modes. Although Quadra and GPT versions have different representation shaping mechanisms, they have their merits and are generally comparable.

**Figure S3**. Identified associations between marker pathways. The distance between connected genes were decided by the significance level of the associations (closer pathways have more significant associations). Names of the pathways can be found in **Table S1**.

**Table S1**. Identified metastasis-related marker genes and pathways.

## References

[1] Z. Hou, V. H.-F. Lee, D. L.-W. Kwong, X. Guan, Z. Liu, and W. Dai, “RNAGAN: Train One and Get Four, Multipurpose Human RNA-Seq Analysis Tool with Enhanced Interpretability and Small Data Size Capability,” bioRxiv, p. 2026.03.17.712527, 2026.

[2] I. Goodfellow et al., “Generative adversarial networks,” Communications of the ACM, vol. 63, no. 11, pp. 139–144, 2020.

[3] M. Marouf et al., “Realistic in silico generation and augmentation of single-cell RNA-seq data using generative adversarial networks,” Nature communications, vol. 11, no. 1, p. 166, 2020.

[4] F. L. Barsha and W. Eberle, “An in-depth review and analysis of mode collapse in generative adversarial networks,” Machine Learning, vol. 114, no. 6, p. 141, 2025.

[5] A. Muneer et al., “From classical machine learning to emerging foundation models: review on multimodal data integration for cancer research,” Artificial Intelligence Review, 2026.

[5] E. Vorontsov et al., ”A foundation model for clinical-grade computational pathology and rare cancers detection,” Nature medicine, vol. 30, no. 10, pp. 2924–2935, 2024.

[7] G. Arango-Argoty et al., ”Pretrained transformers applied to clinical studies improve predictions of treatment efficacy and associated biomarkers,” Nature Communications, vol. 16, no. 1, p. 2101, 2025.

[8] K. A. Tran, O. Kondrashova, A. Bradley, E. D. Williams, J. V. Pearson, and N. Waddell, “Deep learning in cancer diagnosis, prognosis and treatment selection,” Genome medicine, vol. 13, no. 1, p. 152, 2021.

[9] Y. He et al., “Foundation model for advancing healthcare: challenges, opportunities and future directions,” IEEE Reviews in Biomedical Engineering, vol. 18, pp. 172–191, 2024.

[10] L. Zhuang, S. H. Park, S. J. Skates, A. E. Prosper, D. R. Aberle, and W. Hsu, “Advancing precision oncology through modeling of longitudinal and multimodal data,” IEEE Reviews in Biomedical Engineering, 2025.

[11] H. Huang et al., “Immunotherapy for nasopharyngeal carcinoma: current status and prospects,” International journal of oncology, vol. 63, no. 2, p. 97, 2023.

[12] Y.-P. Chen et al., ”Single-cell transcriptomics reveals regulators underlying immune cell diversity and immune subtypes associated with prognosis in nasopharyngeal carcinoma,” Cell research, vol. 30, no. 11, pp. 1024–1042, 2020.

[13] A. D. Colevas et al., “NCCN Guidelines® insights: head and neck cancers, version 2.2025: featured updates to the NCCN guidelines,” Journal of the National Comprehensive Cancer Network, vol. 23, no. 2, pp. 2–11, 2025.

[14] K. C. Wong et al., “Nasopharyngeal carcinoma: an evolving paradigm,” Nature reviews Clinical oncology, vol. 18, no. 11, pp. 679–695, 2021.

[15] X. Li, Y. Guo, M. Xiao, and W. Zhang, ”The immune escape mechanism of nasopharyngeal carcinoma,” The FASEB Journal, vol. 37, no. 7, p. e23055, 2023.

[16] Q. Liu, Y. Tong, T. Sun, L. He, X. Xu, and K. You, “Epstein–Barr virus reprograms immune escape in nasopharyngeal carcinoma,” Virulence, vol. 17, no. 1, p. 2677306, 2026.

[17] H. Liu et al., “Nasopharyngeal carcinoma: current views on the tumor microenvironment’s impact on drug resistance and clinical outcomes,” Molecular cancer, vol. 23, no. 1, p. 20, 2024.

[18] Y. Liu et al., ”Single-cell and spatial transcriptome analyses reveal tertiary lymphoid structures linked to tumour progression and immunotherapy response in nasopharyngeal carcinoma,” Nature Communications, vol. 15, no. 1, p. 7713, 2024.

[19] Z. Hou et al., ”Tertiary lymphoid structure-related RNA indicator as metastasis risk factor in nasopharyngeal carcinoma,” Clinical and Translational Medicine, vol. 15, no. 12, p. e70539, 2025.

[20] C. C. S. Program et al., “CZ CELLxGENE Discover: a single-cell data platform for scalable exploration, analysis and modeling of aggregated data,” Nucleic acids research, vol. 53, no. D1, pp. D886–D900, 2025.

[21] J. Liu et al., “An integrated TCGA pan-cancer clinical data resource to drive high-quality survival outcome analytics,” Cell, vol. 173, no. 2, pp. 400–416. e11, 2018.

[22] R. Edgar, M. Domrachev, and A. E. Lash, “Gene Expression Omnibus: NCBI gene expression and hybridization array data repository,” Nucleic acids research, vol. 30, no. 1, pp. 207–210, 2002.

[23] Y. Liu et al., “Tumour heterogeneity and intercellular networks of nasopharyngeal carcinoma at single cell resolution,” Nature communications, vol. 12, no. 1, p. 741, 2021.

[24] J. L. Ba, J. R. Kiros, and G. E. Hinton, ”Layer normalization,” arXiv preprint arXiv:1607.06450, 2016.

[25] R. Xiong et al., “On layer normalization in the transformer architecture,” in International conference on machine learning, 2020: PMLR, pp. 10524-10533.

[26] L. Huang, Normalization techniques in deep learning. Springer, 2022.

[27] X. Gao et al., ”A comprehensive survey on imbalanced data learning,” Frontiers of Computer Science, vol. 20, no. 11, p. 2011622, 2026.

[28] G. G. Chrysos, S. Moschoglou, G. Bouritsas, Y. Panagakis, J. Deng, and S. Zafeiriou, “P-nets: Deep polynomial neural networks,” in Proceedings of the IEEE/CVF Conference on Computer Vision and Pattern Recognition, 2020, pp. 7325–7335.

[29] The MathWorks Inc. I. The MathWorks. (2024). Deep Learning Toolbox User’s Guide. Available: https://www.mathworks.com/help/deeplearning/

[30] A. Vaswani et al., “Attention is all you need,” Advances in neural information processing systems, vol. 30, 2017.

[31] A. Chowdhery et al., “Palm: Scaling language modeling with pathways,” Journal of machine learning research, vol. 24, no. 240, pp. 1–113, 2023.

[32] S. Yang et al., “Discretizing multiple continuous predictors with U-shaped relationships with ln OR: introducing the recursive gradient scanning method in clinical and epidemiological research,” BMC Medical Research Methodology, vol. 25, no. 1, p. 70, 2025.

[33] Q. Chen, L. Yang, A. Wang, X. Luo, and Y. Zhang, ”QuadEnhancer: Leveraging Quadratic Transformations to Enhance Deep Neural Networks,” Advances in Neural Information Processing Systems, vol. 38, pp. 45730–45749, 2026.

[34] S. Somvanshi, S. Das, S. A. Javed, G. Antariksa, and A. Hossain, ”A survey on deep tabular learning,” arXiv preprint arXiv:2410.12034, 2024.

[35] A. Bose, P. Mukherjee, and A. Sarkar, “GBAT-UNet: group bottleneck transformer with attention in UNet for retinal vessel segmentation,” Neural Computing and Applications, vol. 38, no. 8, p. 274, 2026.

[36] E. A. Eisenhauer et al., “New response evaluation criteria in solid tumours: revised RECIST guideline (version 1.1),” European journal of cancer, vol. 45, no. 2, pp. 228–247, 2009.

[37] J. H. Friedman, T. Hastie, and R. Tibshirani, “Regularization paths for generalized linear models via coordinate descent,” Journal of statistical software, vol. 33, pp. 1–22, 2010.

[38] B. Phipson, S. Lee, I. J. Majewski, W. S. Alexander, and G. K. Smyth, “Robust hyperparameter estimation protects against hypervariable genes and improves power to detect differential expression,” The annals of applied statistics, vol. 10, no. 2, p. 946, 2016.

[39] Y. Li et al., “Multi-omic characterization of nasopharyngeal carcinoma delineates the subtype-specific landscape of response to induction chemotherapy,” Nature Cancer, pp. 1–20, 2026.

