## Supplementary figures and images for "Foundation Model RNAGAN Enhances Biomedical Insight of Nasopharyngeal Carcinoma Metastasis"

### Figure1.tif

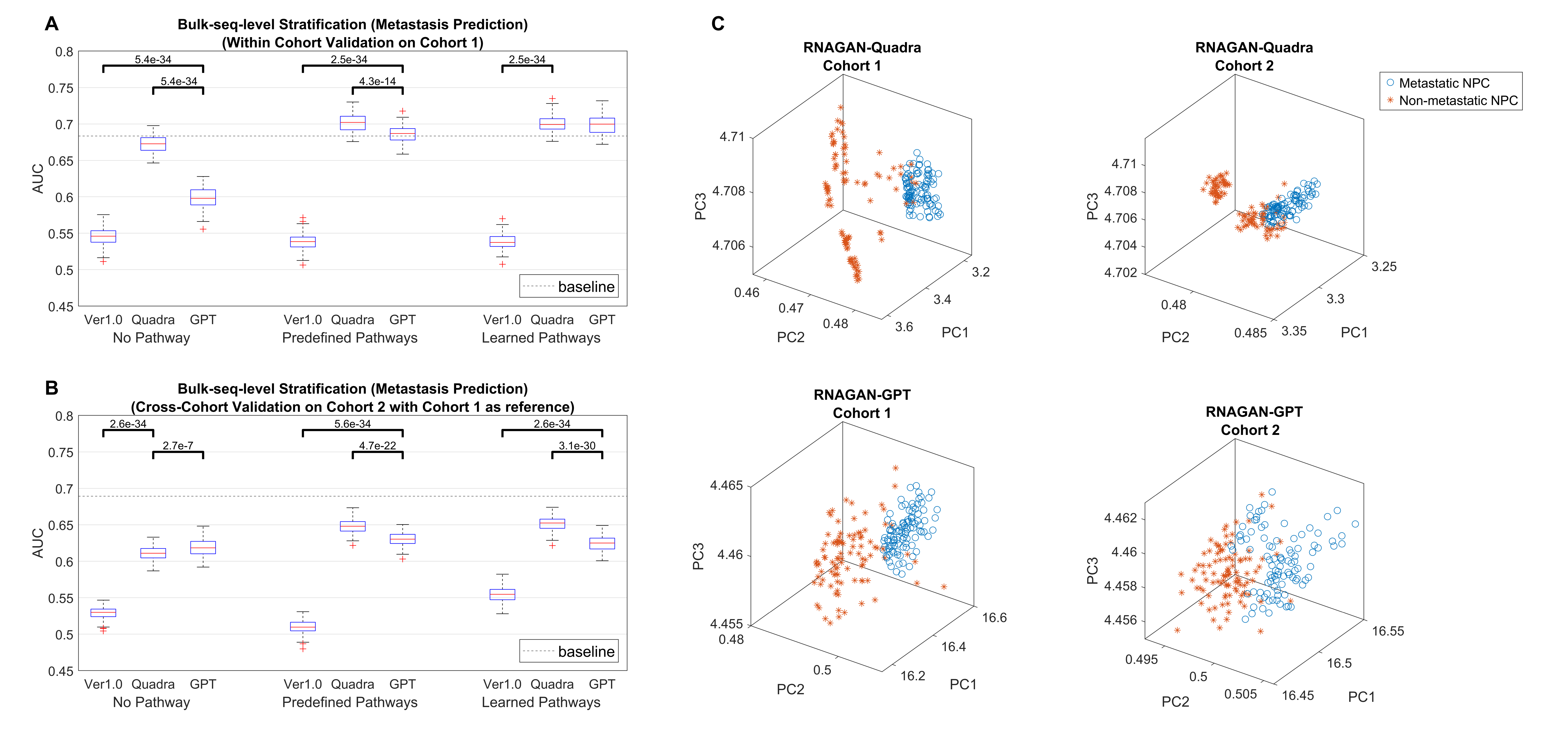

### Figure2.tif

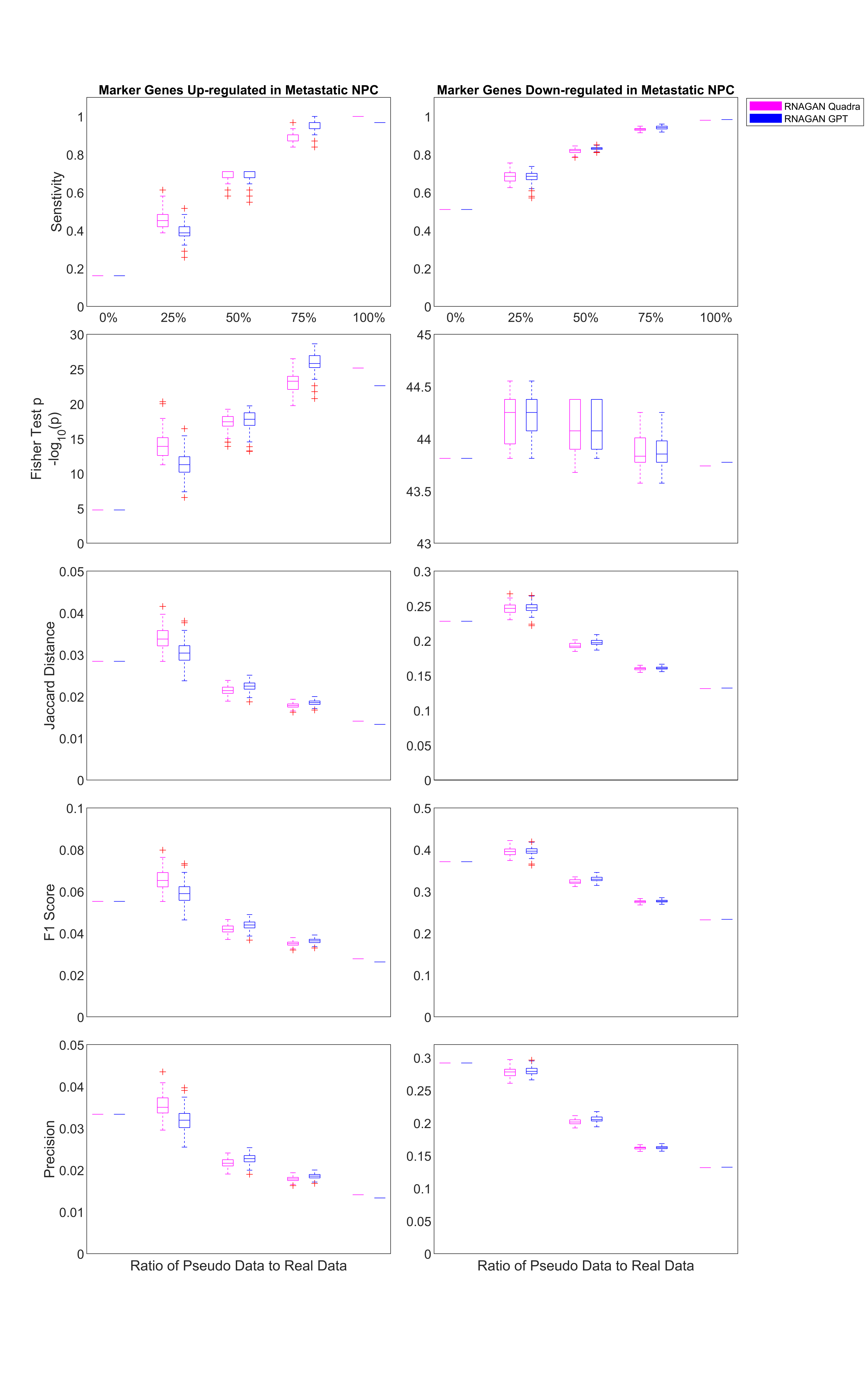

### Figure2C.gif

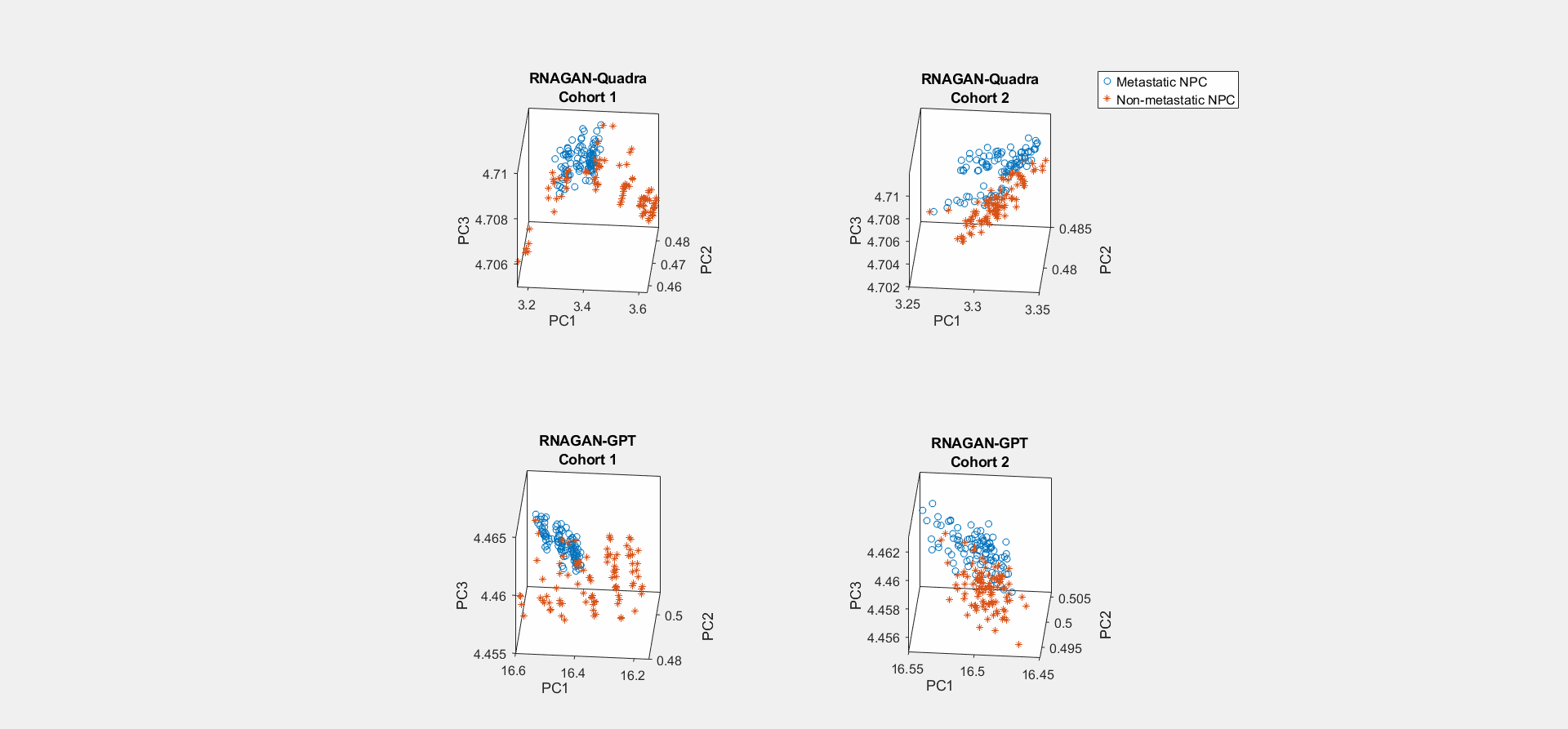

### Figure3.tif

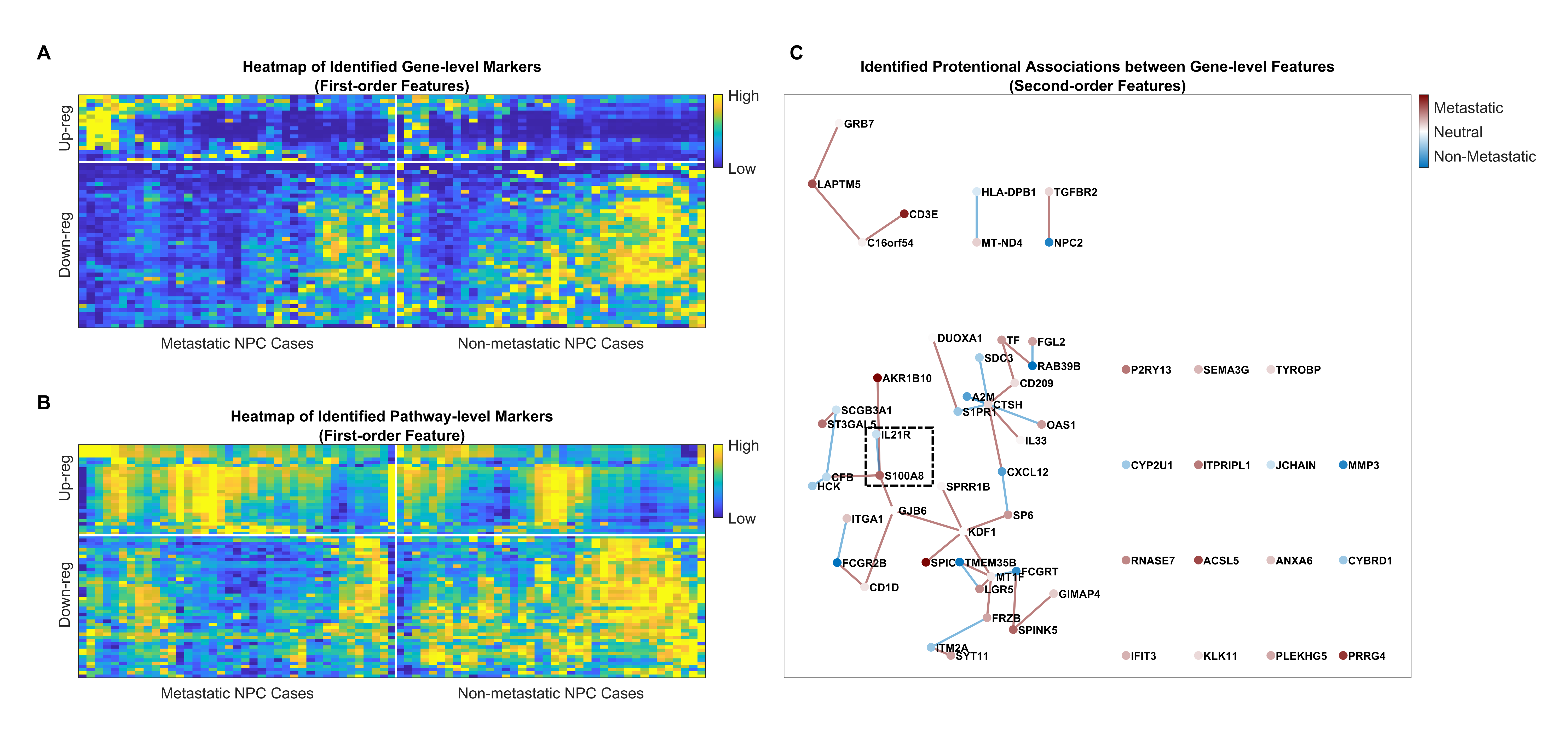

### Figure4.tif

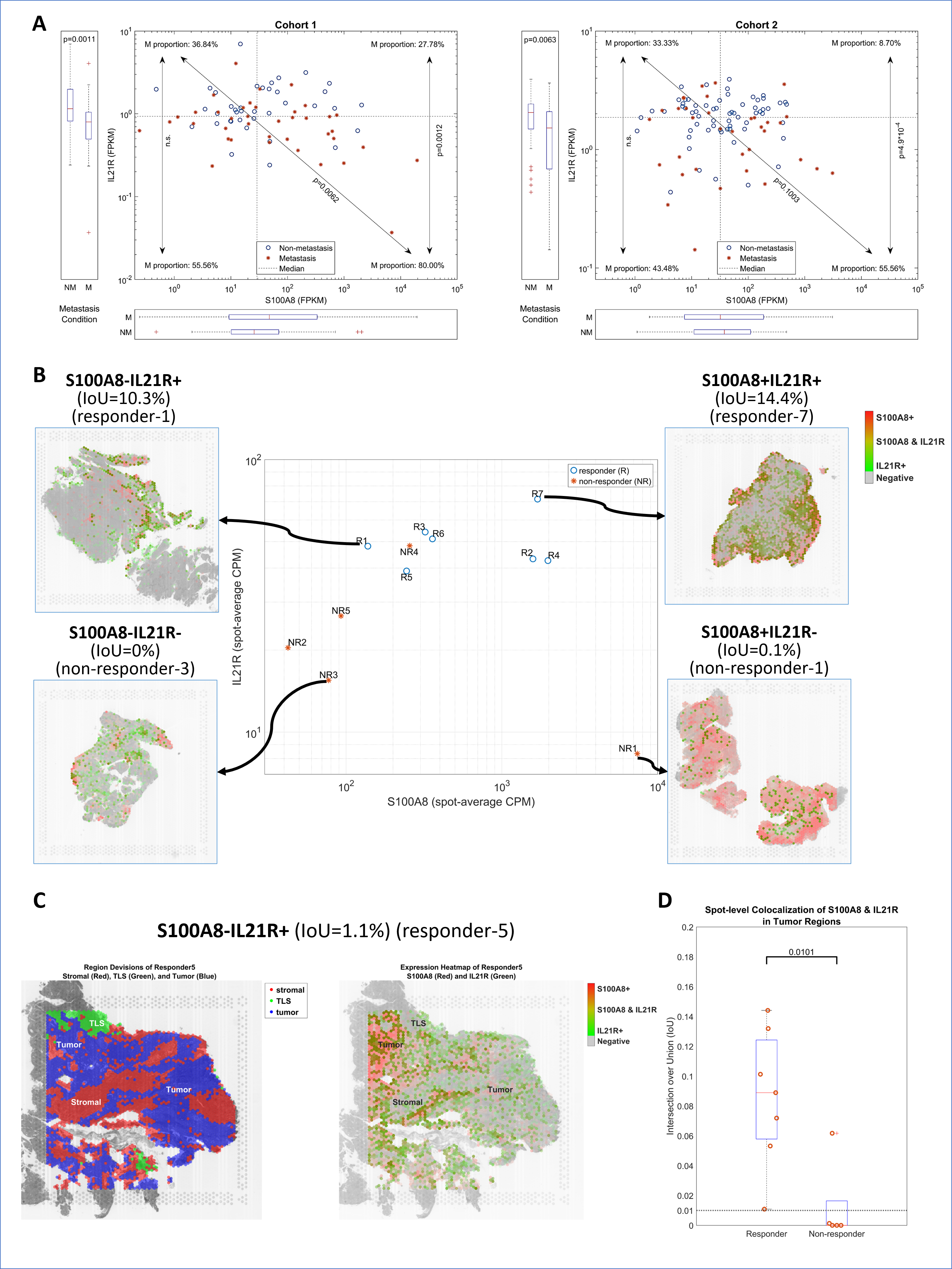

### FigureS1.tif

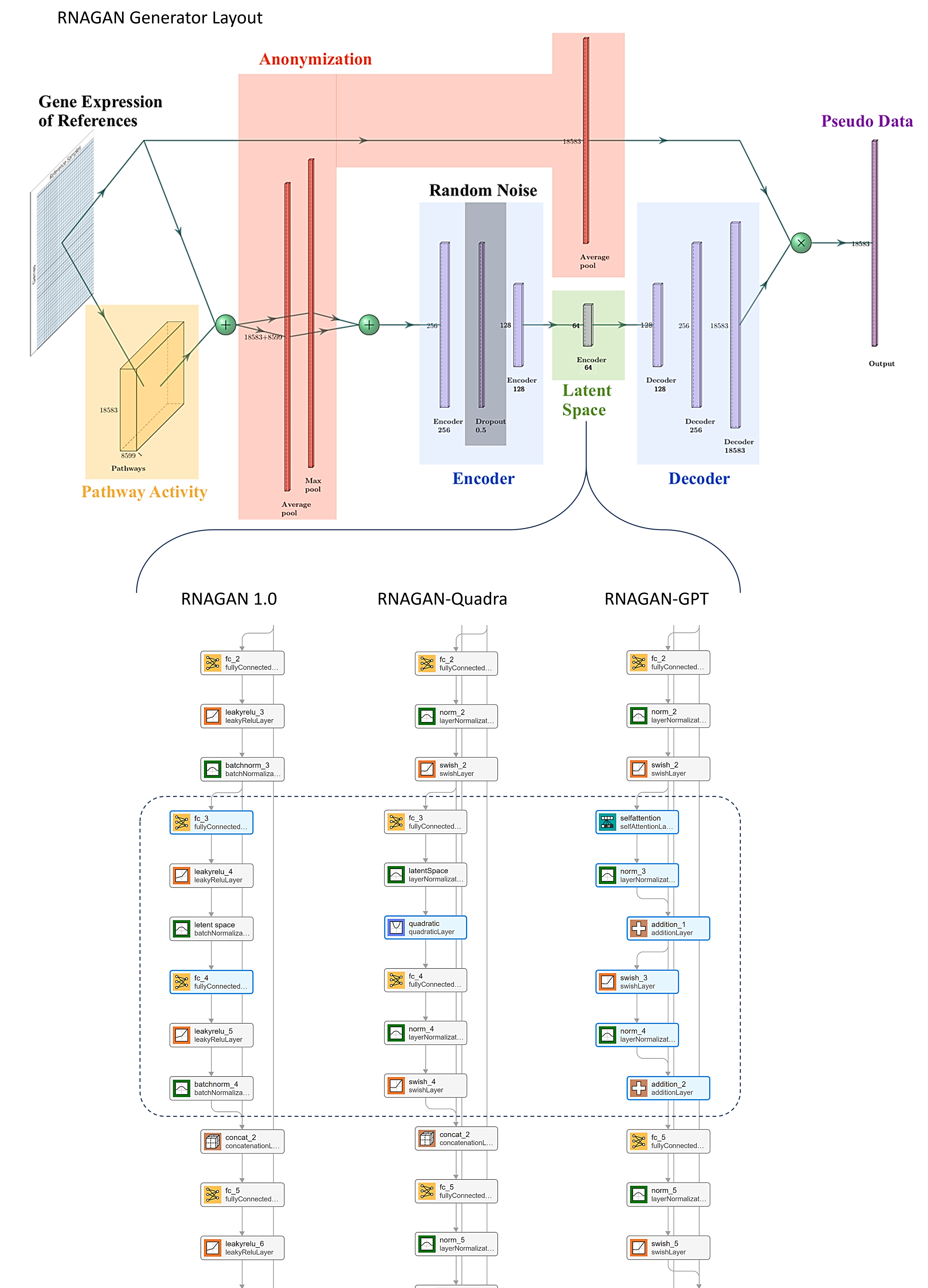

### FigureS2.tif

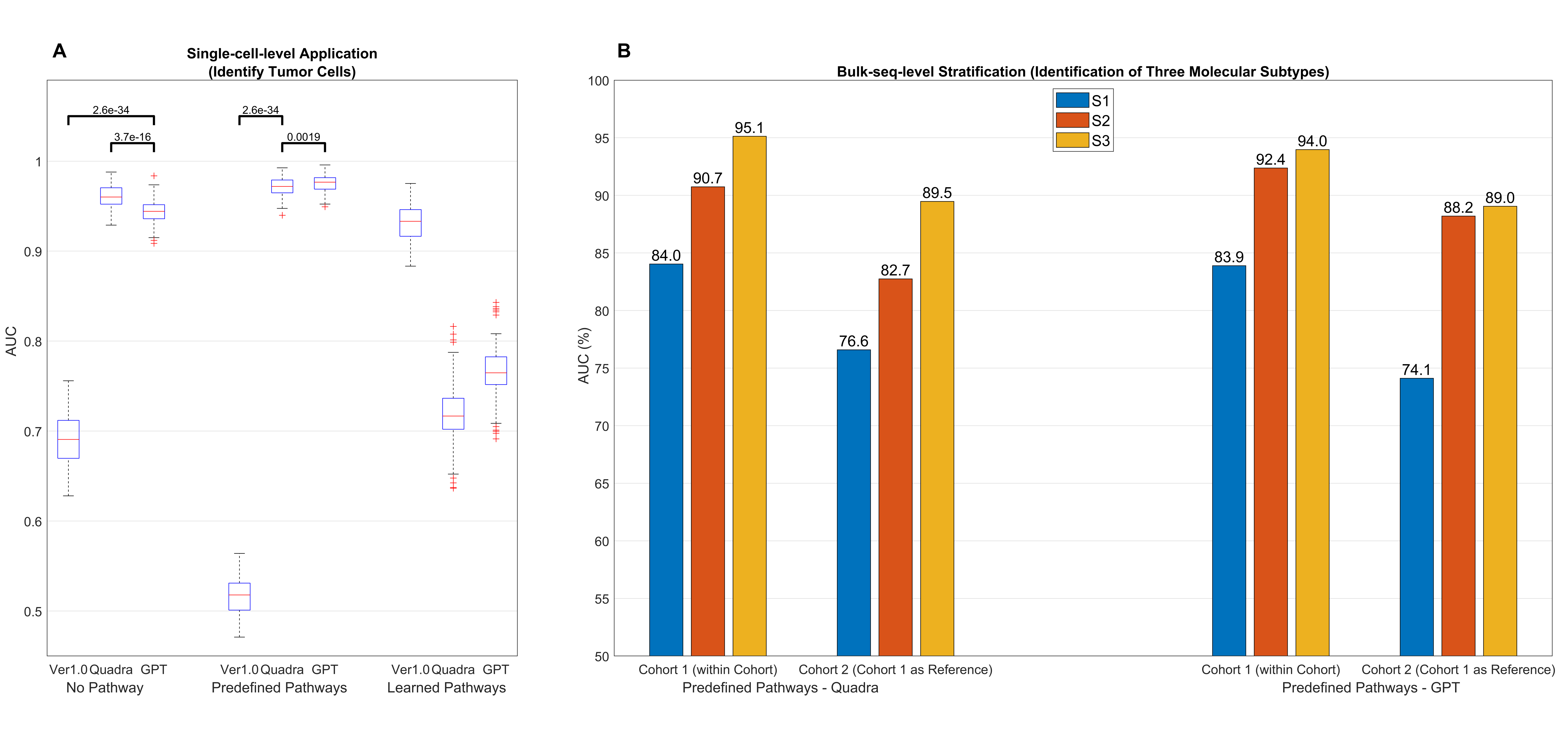

### FigureS3.tif

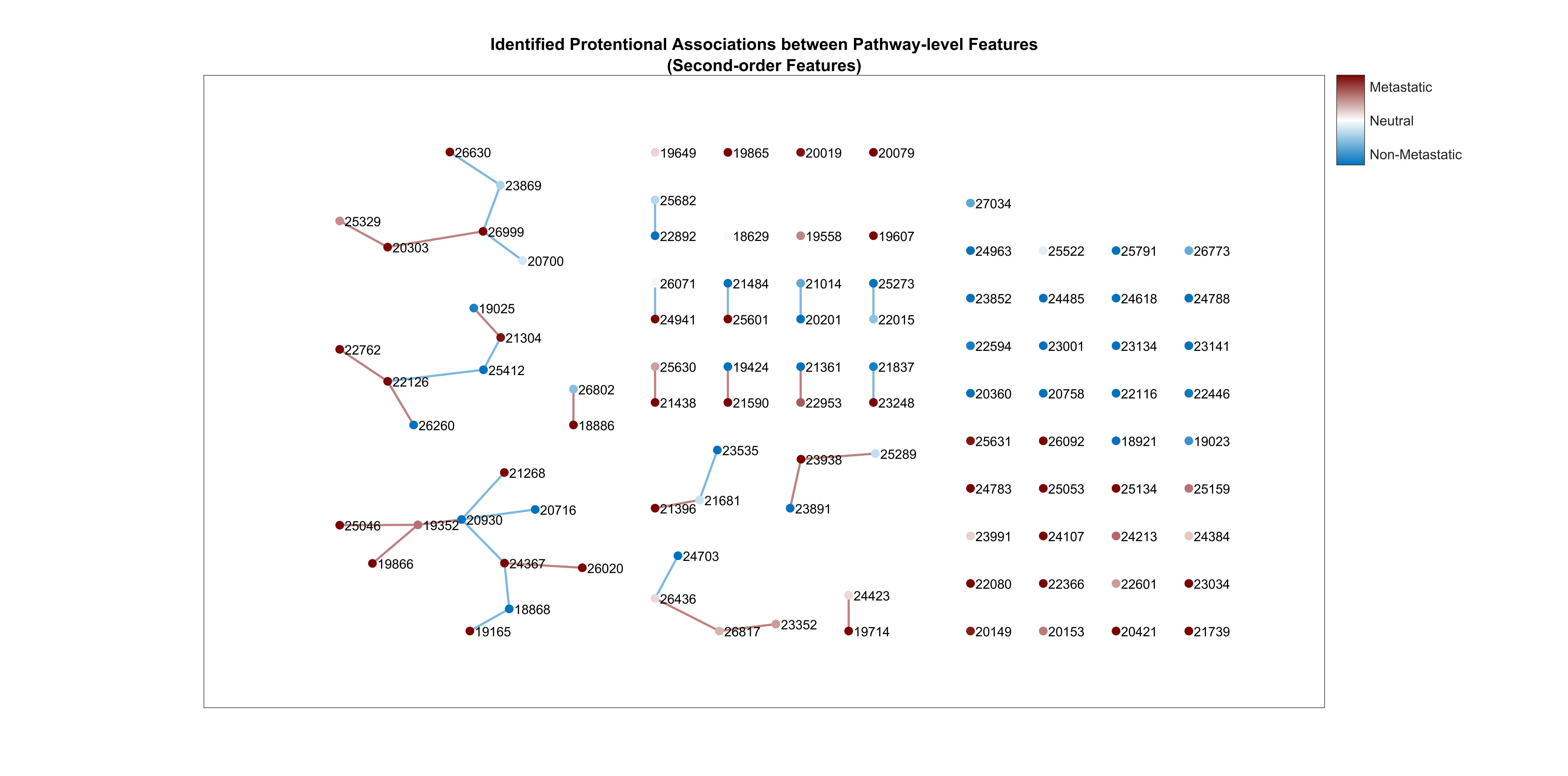
